# Properties of rhizosphere soil associated with herbaceous plant roots analyzed using small-scale protocols

**DOI:** 10.1101/800664

**Authors:** Shinichi Yamazaki, Kumiko Ochiai, Junko Motokawa, Shoichiro Hamamoto, Akifumi Sugiyama, Masaru Kobayashi

## Abstract

The rhizosphere, which is the region of soil adjacent to plant roots, is affected by the activities of both plant roots and associated microorganisms which cause changes in soil properties including nutrient mineral composition. Accordingly, the actual availability of plant nutrients may not be the same as that estimated on the basis of bulk soil analysis. However, the extent and manner in which the availability of plant nutrients in bulk and rhizosphere soils differ remain unclear. Therefore, the present study defined the rhizosphere as the soil adhered to plant roots, established a set of small-scale protocols for analyzing the nutrient minerals of small soil samples, and then characterized the rhizosphere soil of sorghum, *Sorghum bicolo*r (L.) Moench. The mineral contents of the bulk and rhizosphere soil differed significantly, with nutrient contents generally greater in the rhizosphere, and particularly remarkable accumulation was observed in regards to ammonium ion and exchangeable potassium concentrations. Such accumulation might be due, in part, to the greater per weight surface areas of rhizosphere soil particles, but other mechanisms, including the accumulation of organic matter, could also be involved.

## Introduction

The environment in the immediate vicinity of plant roots, the rhizosphere, is generally considered to be distinct from that of the surrounding soil (i.e., bulk soil), and the differences in these two regions can be attributed to multiple factors. Respiration by roots, for example, is associated with the consumption of oxygen and production of carbon dioxide, thereby causing local changes in gas composition. Meanwhile, the excretion of exudates by plant roots provides a food source for heterotrophic microbes and promotes the establishment of distinct microbial communities [1–3]. Furthermore, the selective uptake of nutrients by roots, release of protons or bicarbonate ions in exchange with absorbed nutrients, and the release of plant-derived organic acids or chelators to solubilize sparingly soluble salts cause changes in the chemical composition of the rhizosphere [4]. Together, the resulting differences between the nutrient availabilities of bulk and rhizosphere soil can have significant effects on plant growth. However, the extent and manner in which the availability of plant nutrients in bulk and rhizosphere soils differ remain unclear. For example, the ammonium ion (NH_4_^+^) levels of rhizosphere soil have been reported to be both higher [5–7] or lower [8,9] than that of bulk soil, and the same trend has been reported for potassium (K) [10–12].

To gain insights into the rhizosphere, soil needs to be fractionated according to distance from plant roots. The so-called “rhizobox” technique, for example, allows roots grow within a compartment of soil that is separated from the surrounding region with fine mesh, which prevents the roots from mixing with the surrounding soil but allows the influx and efflux of nutrients and root exudates, respectively. When using this technique, the surrounding soils are fractionated according to their distance from the root-containing compartment, and are analyzed for a variety of physiochemical characteristics, including mineral contents. However, even though the rhizobox technique clearly facilitates the collection and analysis of soil that is free of root contamination, it still fails to provide a complete picture of the rhizosphere since the soil recovered is not actually in direct contact with the roots.

Another approach used to study rhizosphere soil is the collection of adhered soil from root surfaces by gentle brushing, after the removal of most of the soil trapped in the root system. Even though it is possible that soil collected in this manner can be contaminated with root fragments, the soil samples can provide unique insight into the environmental conditions of the immediate vicinity of the roots. As such, the method has been used for a variety of purposes, such as analyzing the structures of rhizosphere microbial communities [13,14] and investigating the nutrient availability and dynamics of tree-associated rhizospheres [7,15]. However, little information is available regarding the application of this method to investigate the nutrient availability in the rhizosphere of herbaceous plant species, probably because of the difficulty in recovering enough root-adhering soil for the analysis. Therefore, establishing methods for analyzing the chemical properties of small soil samples would likely facilitate research into the nutrient dynamics of rhizosphere soil associated with agriculturally important herbaceous crop species.

Accordingly, the present study defined the rhizosphere as the soil adhered to plant roots, established a set of small-scale protocols for analyzing the nutrient contents of small soil samples, and characterized the rhizosphere soil of sorghum, *Sorghum bicolo*r (L.) Moench.

## Materials and methods

### Plant cultivation and soil collection

For plant cultivation, soil was collected from an experimental farm on the campus of Kyoto University, mixed with an equal volume of commercial potting soil (Tachikawa Heiwa Noen, Tochigi, Japan), and passed through a 5-mm sieve to remove large aggregates. Portions of the mixed soil (10 kg) were then transferred to 1/2000-are pots, which were placed in a greenhouse, and fertilized with urea, KH_2_PO_4_, and KCl at 100 kg N ha^−1^, 100 kg P_2_O_5_ ha^−1^ and 100 kg K_2_O ha^−1^. Nine pots were prepared so that the soil could be sampled at three time points, with three replicates per sampling time. Three seeds (*Sorghum bicolor* L. Moench ‘BT×623’) were sown in each pot, and at 2 weeks after germination, the plants were thinned to one plant per pot.

Rhizosphere soil samples were collected at 6, 11, and 16 weeks after germination, which corresponded to the vegetative, heading, and harvest stages of the sorghum plants, respectively. During each scheduled collection, the roots were gently removed from the soil and shaken vigorously by hand, and the soil adhered to the roots was gently brushed off using a clean paint brush and collected. Meanwhile, bulk soil samples were collected from the soils remaining in the pots. Both sample types (bulk and rhizosphere) were air-dried at room temperature and passed through a 500-µm sieve. In the experiments examining the effects of particle size on NH_4_^+^ contents, the samples were further fractionated by passing through a 200-µm sieve. The fraction that did not pass through the sieve was designated as coarse fraction (200-500 µm), whereas that did pass was as fine fraction (<200 µm).

Bulk and rhizosphere soil samples were also collected from wild sedge (*Cyperus* spp.) growing in the uncultivated portion of the experimental farm in Kyoto University, as described above for sorghum.

### Total carbon and nitrogen measurement

The total carbon (C) and nitrogen (N) contents of the soil samples were measured using 40 mg of each sample and an NC analyzer (Sumigraph NC-22F; Sumika Chemical Analysis Service, Ltd., Osaka, Japan).

### Inorganic nitrogen measurement

Inorganic N was extracted from each sample by suspending 30 mg soil in 300 µL 0.5 M K_2_SO_4_ and shaking the suspension at 150 rpm at room temperature for 1 h. Then, after centrifuging the suspension at 13,000 ×*g* for 10 min, the resulting supernatant was used for the measurement of ammonium (NH_4_^+^) and nitrate (NO_3_^−^) contents.

The NH_4_^+^ contents were measured using a simplified indophenol blue method [16] that was modified to accommodate a microplate format. In the wells of a 96-well microplate, 50 µL extract or standard solution (0–250 ng NH_4_-N as ammonium sulfate) was mixed with 20 µL of a solution that contained potassium sodium tartrate (0.212 M), trisodium citrate (0.136 M), and HCl (4 mM) and then 40 µL of freshly prepared 2:1:3 (v:v:v) mixture of boric acid-NaOH (20 mM and 0.4 M, respectively), 2-phenylphenol sodium salt (0.313 M), and sodium pentacyanonitrosylferrate (III, 1.0 mM). The stock solutions of 2-phenylphenol sodium salt and sodium pentacyanonitrosylferrate (III) were stored refrigerated in the dark. Blue color was developed by adding 90 µL of a sodium hypochlorite solution that contained 0.025–0.040% active chlorine, covering the plate with plastic wrap, and incubating the plate at 37°C for 20 min. The absorbances (655 nm) of the samples were then measured using an SH-1200Lab microplate reader (Corona Electric, Ibaraki, Japan).

Meanwhile, the NO_3_^−^ contents were measured using a modified version of the Cataldo method [17]. Aliquots (50 μL) of extract or standard solution (0–400 ng NO_3_-N as KNO_3_) were added to the wells of a 96-well microplate and dried in an oven at 70°C. The resulting crystals were covered with 10 µL salicylic acid (0.05 g mL^−1^ in sulfuric acid) and incubated at 80°C for 20 min. Then, 250 μL NaOH (2 M) was added to each well, and the crystals were dissolved completely by pipetting. After incubation for 20 min at room temperature, the absorbances (410 nm) of the samples were measured using an SH-1200Lab microplate reader.

### Available nitrogen estimation

The available N contents of the soil samples were estimated using neutral phosphate buffer extraction [18]. Fifty milligrams of each soil sample was suspended in 250 µL neutral phosphate buffer, which was prepared as a 35:65 (v:v) mixture of KH_2_PO_4_ (66.7 mM) and Na_2_HPO_4_ (66.7 mM), shaken at room temperature for 1 h at 150 rpm, and centrifuged at 13,000 ×*g* for 10 min. Aliquots (200 µL) of each supernatant were transferred to a microplate, and absorbance (420 nm) was measured using an SH-1200Lab microplate reader. Since the reported method [18] uses absorbance values that are measured using a 1-cm optical path length (*A*′_420_), the measured absorbance values (*A*_420_) were converted using the following empirically determined formula:

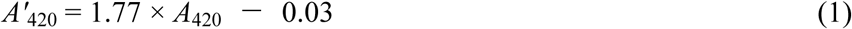

The phosphate buffer-extractable and available N contents (mg N kg^−1^) were then estimated using the following equations [19]:

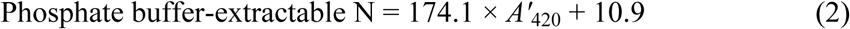

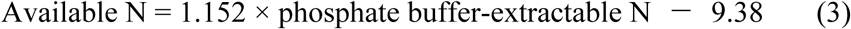

### Available phosphorus measurement

The available phosphorus (P) contents of the soil samples were estimated using the Truog method [20]. Twenty milligrams of each soil sample was suspended in 4 mL H_2_SO_4_ (1 mM) that contained 0.3% (w/v) ammonium sulfate, in a 5-mL plastic tube, and shaken at room temperature for 30 min. A 1-mL aliquot of each suspension was then transferred to a 1.5-mL microtube and centrifuged at 13,000 ×*g* for 10 min. Aliquots (200 µL) of the resulting supernatants or standard solutions (0–600 ng P_2_O_5_ as KH_2_PO_4_) were then transferred to a microplate, mixed with 50 µL coloring reagent, and incubated at room temperature for 20 min. The coloring reagent was prepared fresh on the day of use by formulating a 10:3:1:11 (v:v:v:v) mixture of H_2_SO_4_ (2.5 M), ammonium molybdate (4%, w/v), potassium antimonyl tartrate trihydrate (0.28%, w/v), and distilled water, and then dissolving ascorbic acid into the mixture at a concentration of 4.2 mg mL ^−1^. The absorbances (880 nm) of the prepared sample reactions were measured using an SH-1200Lab microplate reader.

### Exchangeable base measurement

Exchangeable bases were extracted from each soil sample by suspending 30 mg soil in 600 µL ammonium acetate (1 M, pH 7.0) and shaking the suspension at 150 rpm at room temperature for 1 h. Then, after being centrifuged at 13,000 ×*g* for 10 min, the resulting supernatant was diluted five-fold with distilled water and used for the measurement of K by flame photometry (AA-6200; Shimadzu, Kyoto, Japan). Portions of each diluted supernatant were further diluted with distilled water, mixed with 1/100 volumes of LaCl_3_ (0.4 M), which functioned as an interference suppressor, and used for the measurement of calcium (Ca) and magnesium (Mg) using an AA-6200 atomic absorption spectrometer. Meanwhile, the residue from the base extraction with ammonium acetate was mixed with 1 mL ethanol (80%, v/v) by vortexing and centrifuged at 13,000 ×*g* for 10 min to remove the supernatant. The operation was repeated two times, and resulting precipitate was resuspended in 600 µL KCl (2 M) and shaken for 1 h at 150 rpm at room temperature. Then, after centrifuging at 13,000 ×*g* for 10 min, the resulting supernatant was subjected to the simplified indophenol blue method, as described above, in order to quantify the NH_4_^+^ content and, thus, the cation exchange capacity (CEC) of the soil. Finally, base saturation was calculated as the ratio of the sum of exchangeable K, Ca, and Mg to CEC.

### Standard soil analysis

Bulk soil samples that were collected from various farmers’ fields in Kyoto were analyzed using both the small-scale and standard protocols. The analysis using standard protocols was conducted by Seikaken, Inc. (Kumamoto, Japan).

### Soil particle analysis

To measure particle size distribution, the organic matters in each soil sample were first removed by mixing dry samples (~10 g for bulk soil and ~0.5 g for rhizosphere soil) with ~200 mL hydrogen peroxide (6%, w/v). Each resulting soil suspension was filtered through a 200-μm sieve, mixed with sodium hexametaphosphate (0.07 M), as a dispersing agent, shaken for 1 d at 25°C, and subject to the measurement of particle size distribution using a laser-diffraction particle size analyzer (SALD-7500nano, Shimadzu). According to the International Society of Soil Science (ISSS) standard [21], <2-µm and 2-to 20-µm particles were classified as clay and silt, respectively. The remaining 20- to 200-µm particles were classified as sand. This analysis was performed with a limited number of samples from the heading or harvest stage (4 samples for bulk and 3 for rhizosphere soil, as indicated in the Supporting information S3) since the amount of soil recovered from the vegetative stage plants were not sufficient for the analysis.

The nitrogen (N_2_) adsorption surface areas of dry bulk and rhizosphere soil samples (0.5–1.0 g) were measured, in triplicate, on the basis of the Brunauer-Emmett-Teller theory [22] using a surface area and porosity analyzer (TriStar II series; Micromeritics Instrument Co., Norcross, GA, USA). The analysis was performed with the soil samples of harvest stage (Supporting information S3).

## Results and discussion

### Protocols for small-scale soil analysis

The aim of the present study was to analyze the chemical properties of the soil directly attached to the surfaces of herbaceous plant roots. However, these methods yielded relatively small soil samples. For example, only 0.4–3 g air-dried soil was obtained from individual container-grown, ~60-cm-tall sorghum plants, and such amounts are insufficient for analyzing soil chemical properties using standard protocols. Therefore, a set of small-scale protocols was developed for the measurement of soil fertility using small amounts of soil. Most of the protocols were developed by simply scaling-down standard protocols. Available N, which is the fraction of soil organic N that is mineralizable during plant growth, is often estimated from the amount of inorganic N that is released during the incubation of soils under specific temperature and moisture conditions [23]. However, because the method is time-consuming and difficult to conduct using small amounts of soils, the present study utilized a simplified method that relies on the phosphate buffer extraction of soil organic N, the method which also has been used frequently to estimate soil available N [24].

To confirm the validity of measurements using the small-scale protocols, a set of soil samples were analyzed using both small-scale and standard protocols, and the resulting values were compared. Since the outsourced analysis using standard protocols did not include the measurement of total C, total N, and available N, only CEC and the inorganic N, available P, and exchangeable cation contents were compared. As shown in Fig 1, the measurements of the small-scale and standard protocols were generally well correlated, and the regression slopes for each parameter was close to 1. These results validate the use of small-scale protocols for the analysis of soil chemical properties from small samples. The amount required for the entire set of analyses was ~200 mg per replicate, which should even be practical to obtain from the roots of herbaceous plants. It is important to note that such small-scale analysis could be biased during the sampling of heterogeneous soils. However, there was no concern about such bias in the present study because the soil samples were passed through a 500-µm sieve to remove large aggregates and could be easily homogenized thereafter.

**Fig 1.**
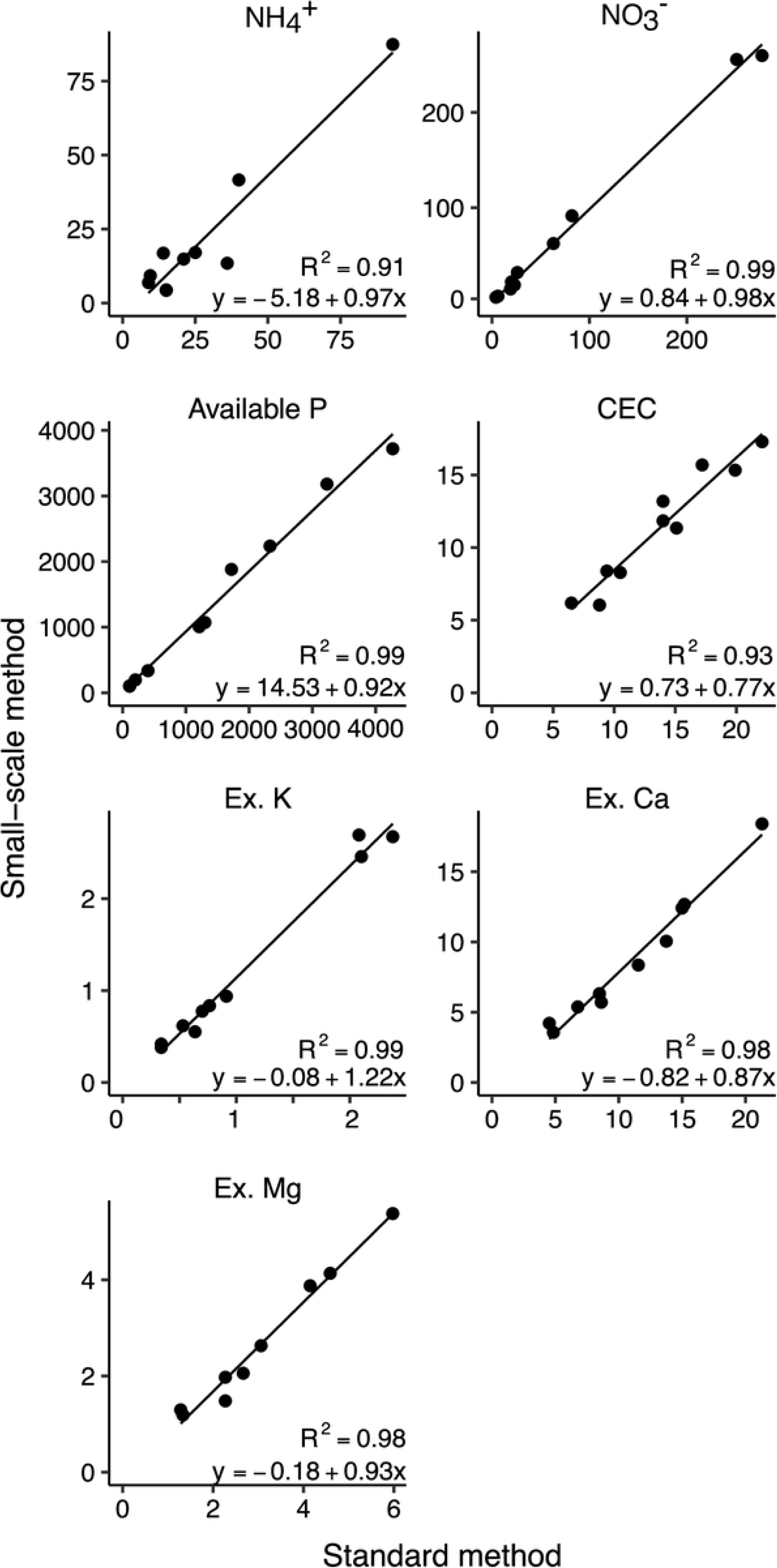
Comparison of standard and small-scale measurements of soil fertility. Bulk soil samples taken from farmers’ fields in Kyoto were analyzed using standard and small-scale methods. Units of values: NH_4_^+^ and NO_3_^−^, mg N kg^−1^ soil; available P, mg P_2_O_5_ kg^−1^ soil; CEC and exchangeable bases (Ex. K, Ex. Ca, and Ex. Mg), cmol_c_ kg^−1^ soil.

### Rhizosphere mineral contents

The bulk and rhizosphere soils of the container-grown sorghum plants were analyzed using the small-scale protocols in order to determine how the soils differed in terms of plant nutrient mineral contents. Most of the parameters were greater in the rhizosphere soil than in the bulk soil (Fig 2). However, neither available P nor base saturation differed significantly between the soil types, and because total C and total N increased concomitantly, the C/N ratios of the bulk and rhizosphere soils were similar (Fig 2).

**Fig 2.**
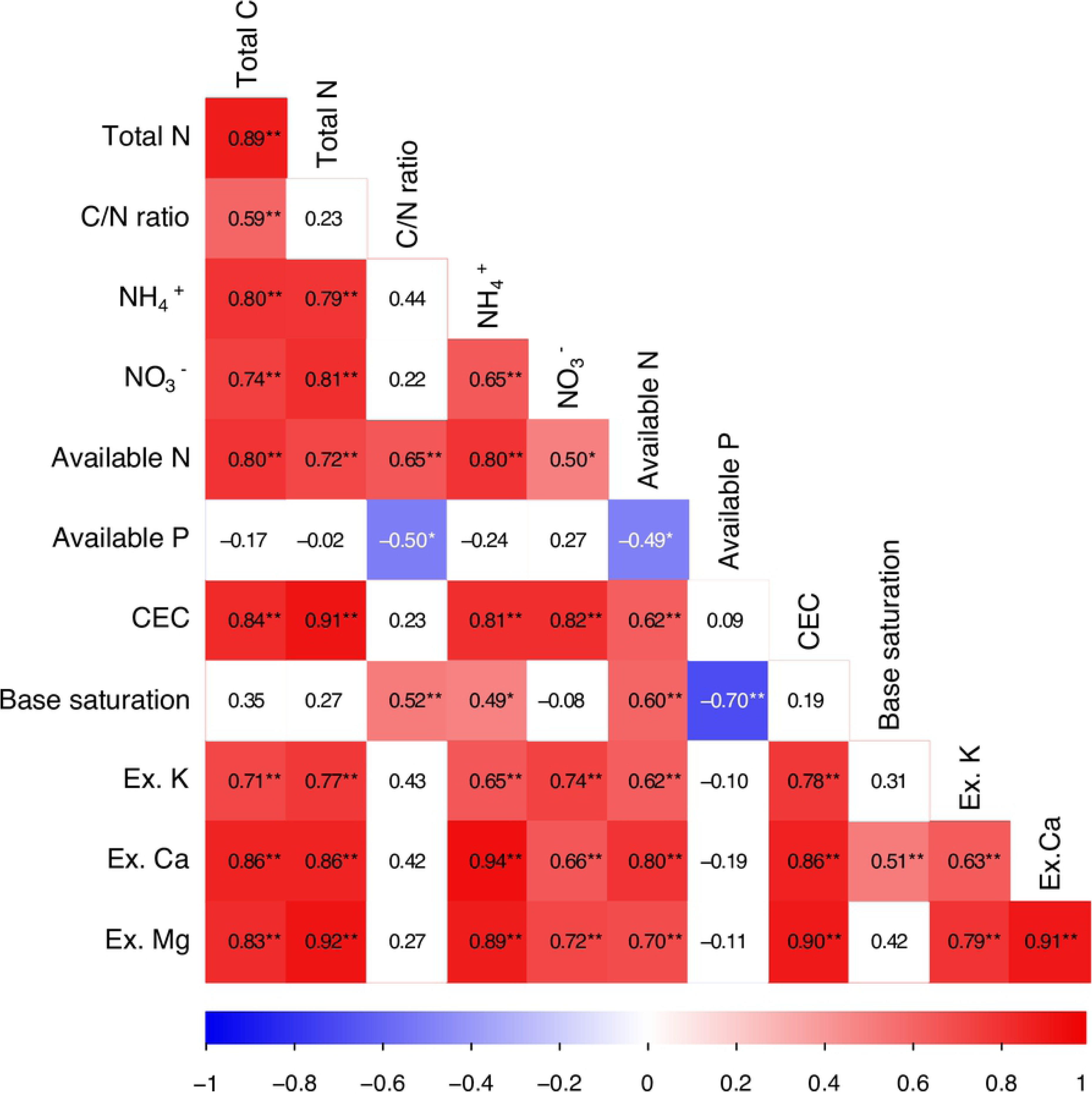
Nutrient mineral contents of bulk and rhizosphere soils associated with container-grown sorghum. CEC, cation exchange capacity; Ex. K, exchangeable K; Ex. Ca, exchangeable Ca; Ex. Mg, exchangeable Mg. Values and error bars represent means ± SE (n=3). Asterisks (*) indicate significant differences between the bulk and rhizosphere soils, according to Student’s T-test (*p* < 0.05). Six, 11, and 16 weeks after germination corresponded to vegetative, heading, and harvest stage, respectively

To determine whether similar differences could also be observed in bulk and rhizosphere soils associated with other plant species, the soils around the roots of wild-growing sedges (*Cyperus* spp.) were also investigated. As in sorghum, greater levels of mineral nutrients were observed in the rhizosphere soil than in the bulk soil (Fig 3), and concomitantly increasing total C and total N contents resulted in similar bulk and rhizosphere soil C/N ratios. The rhizosphere soil of the sedges contained much less NO_3_^−^ than that of the sorghum, probably because the sedges grew wild and were not fertilized. However, the NH_4_^+^ contents of the sorghum and sedge rhizospheres were relatively similar (Figs 2 and 3).

**Fig 3.**
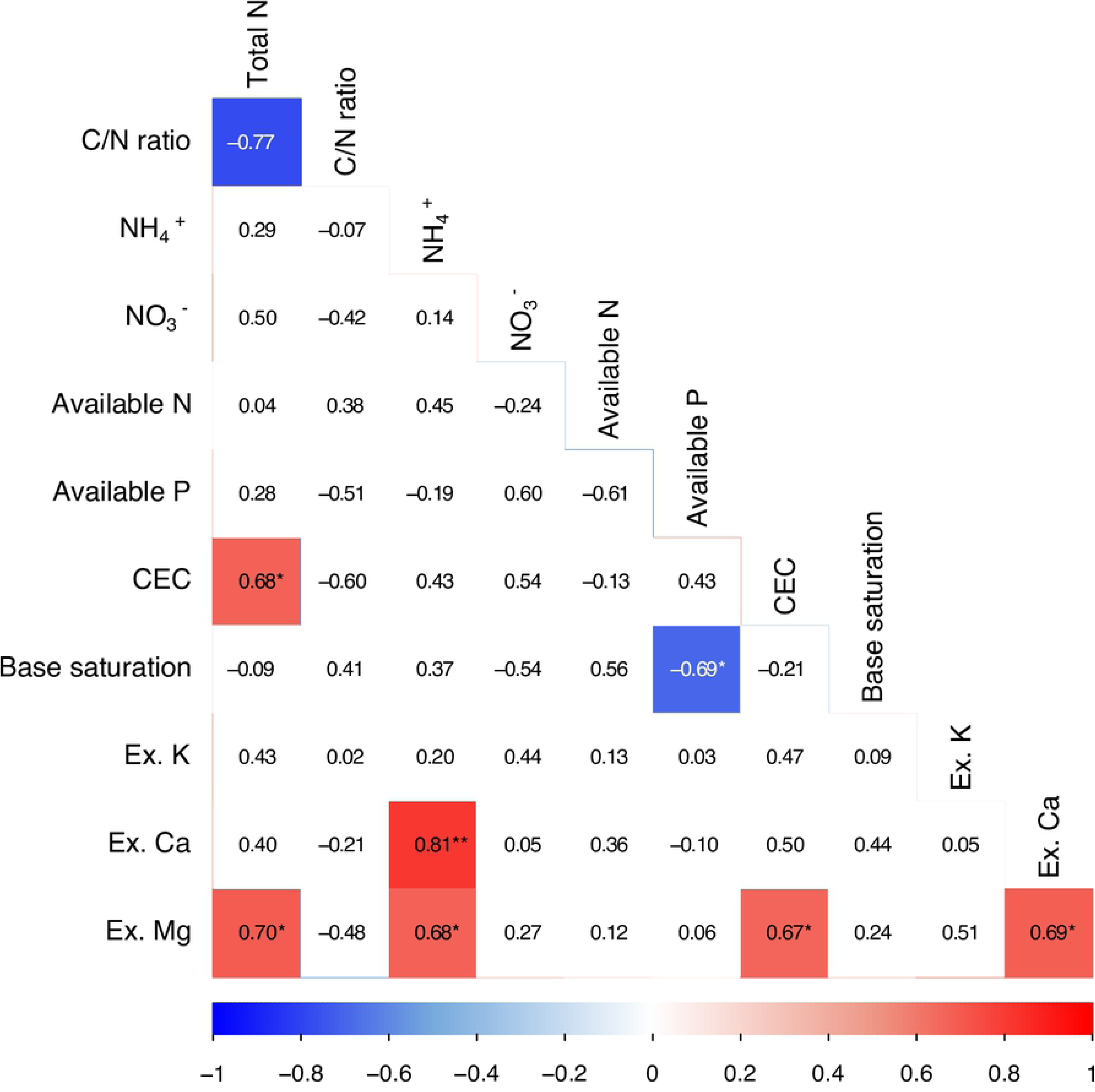
Nutrient mineral contents of bulk and rhizosphere soil associated with wild-growing sedge. Abbreviations are as shown in Fig 2. Values and error bars represent means ± SE (n=3). Asterisks (*) indicate significant differences between the bulk and rhizosphere soils, according to Student’s T-test (*p* < 0.05).

Together, these results demonstrate that the mineral contents of the bulk and rhizosphere soils associated with both sorghum and sedges were distinct, with significantly greater total C, total N, NH_4_^+^, and exchangeable base contents in the rhizosphere than in the surrounding soil (Figs 2 and 3). Indeed, while previous studies of woody species have reported that the mineral contents of rhizosphere soils are generally greater than those of bulk soils [5–7], the results of the present study suggest that such enrichment occurs in annual herbaceous plant species as well.

### Different mineral contents of bulk and rhizosphere soils

Another aim of the present study was to elucidate the mechanisms underlying observed differences in the mineral contents of bulk and rhizosphere soils. Because the rhizosphere soils analyzed in the present study were collected by brushing roots, it is possible that the samples contained fragments of root tissues. However, since inorganic NH_4_^+^ is rarely accumulated by plant cells, the observed enrichment of NH_4_^+^ (Figs 2 and 3) cannot be ascribed to contamination by fine roots.

It is also possible that the observed differences in mineral contents were due to differences in the particle sizes of the samples. Because all the samples were passed through a 500-µm sieve, both the bulk and rhizosphere soil consisted of particles that were less than 500 µm in size. However, since the rhizosphere soil samples only included soil particles that remained adhered to the roots after shaking, the samples may have been enriched with lighter, smaller particles. Since the surfaces of soil particles serve as sites for the adsorption of materials including plant nutrient minerals, smaller particles possess greater surface areas on a per weight basis, thus, more mineral adsorption sites. Therefore, the enrichment of smaller soil particles, if any, could increase the mineral contents of soils on a per weight basis. Accordingly, the particle size distribution of bulk and rhizosphere soils associated with the container-grown sorghum were analyzed. As shown in Table 1, both fractions were rather sandy, and the percentage of clay (<2-µm particles) in each soil type was similar. However, the percentage of silt (2-to 20-µm particles) was greater in the rhizosphere soils than in the bulk soils (Table 1), which indicated that the rhizosphere soil was, indeed, enriched with smaller particles. Consistent with this estimation, the rhizosphere soils had, on average, 30% greater specific surface area than the bulk soils (Table 2). These results suggest that a higher number of adsorption sites per unit weight of soil might account for the greater mineral contents of the rhizosphere soils. This may be especially true for cations, which can attach to negatively charged soil particles through electrostatic interactions. However, the differences in the NH_4_^+^ and K^+^ contents of the bulk and rhizosphere soils were so remarkable that it seems unlikely that the differences are the result of differences in specific surface area alone (Table 2).

**Table 1.** Particle size distribution of bulk and rhizosphere soils associated with container-grown sorghum

|  | Composition (%)<sup>1,2</sup> |  |  |
| --- | --- | --- | --- |
|  | Sand | Silt | Clay |
| Bulk | 73.1 ± 3.81a | 20.4 ± 2.32a | 6.6 ± 1.60a |
| Rhizosphere | 65.5 ± 2.77b | 27.3 ± 2.17b | 7.2 ± 0.70a |

<sup>1</sup> The terms sand, silt, and clay refer to >20-μm, 2- to 20-μm, and <2-μm particles, respectively.
<sup>2</sup> Values represent means ± SD (n=3). Different lowercase letters indicate significant differences between bulk and rhizosphere soils, according to Student's T-test ( $p < 0.05$ ).

**Table 2.**
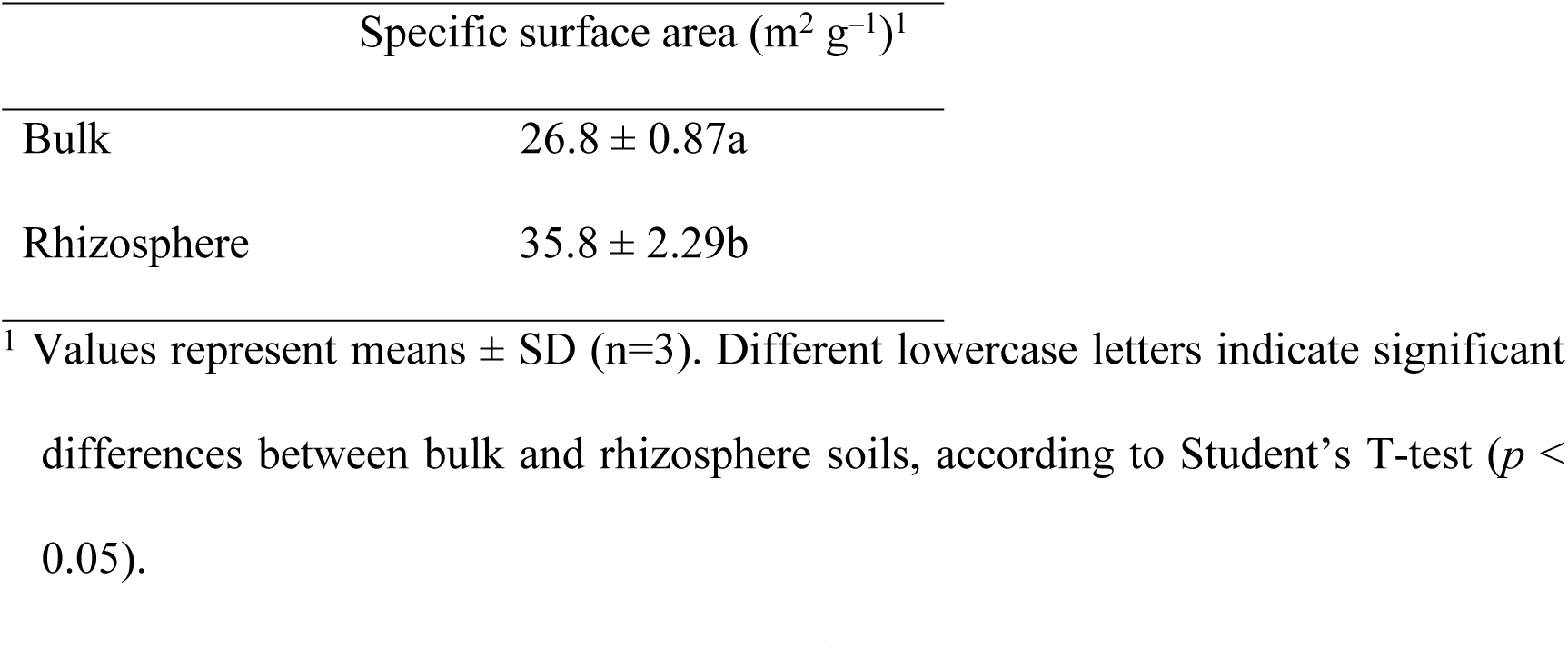
Specific surface areas of bulk and rhizosphere soils associated with container-grown sorghum

| | Specific surface area ($\text{m}^2 \text{g}^{-1}$)<sup>1</sup> |
| --- | --- |
| Bulk | 26.8 ± 0.87a |
| Rhizosphere | 35.8 ± 2.29b |

<sup>1</sup> Values represent means ± SD (n=3). Different lowercase letters indicate significant differences between bulk and rhizosphere soils, according to Student's T-test ( $p < 0.05$ ).

Interestingly, when the rhizosphere soil was separated into coarse (200–500 µm) and fine (<200 µm) particle fractions using a 200-µm sieve, the NH_4_^+^ contents of the fractions were not significantly different from that of unfractionated rhizosphere soil (Table 3). Because NH_4_^+^ is considered to be retained on silt and clay particles (<200 µm), it was assumed that the coarse fraction (200–500 µm) would contain less NH_4_^+^. However, this was not the case, which suggests that silt and clay particles in the rhizosphere were retained in both the fine and coarse soil fractions, probably as components of soil aggregates.

**Table 3.** Ammonium contents of size-fractionated rhizosphere soils associated with container-grown sorghum

| Fraction<sup>1</sup> | $\text{NH}_4^+\text{-N}$ content ($\text{mg kg}^{-1}$ soil)<sup>2</sup> |
| --- | --- |
| Unfractionated | $10.6 \pm 4.67$ |
| Coarse (200–500 $\mu\text{m}$) | $14.9 \pm 6.98$ |
| Fine ($<200 \mu\text{m}$) | $10.9 \pm 4.67$ |

<sup>1</sup> Unfractionated, rhizosphere soil samples taken from the heading-stage plants; Coarse, particles that did not pass through 200- $\mu\text{m}$ sieve; Fine, particles that did pass through 200- $\mu\text{m}$ sieve.
<sup>2</sup> Values represent means ± SD ( $n=3$ ). Values did not vary significantly, according to one-way ANOVA ( $p < 0.05$ ).

To gain further insights into the mechanisms underlying the observed mineral enrichment of the rhizosphere, correlation analysis was performed for the parameters measured from both the bulk and rhizosphere soils associated with container-grown sorghum. Because some of the parameters were not normally distributed owing to differences between the bulk and rhizosphere soils, the Spearman’s rank coefficient was used. As shown in Fig 4, many pairs of parameters yielded high correlation coefficients, which suggested that the parameters were not independent and were indirectly correlated through relationships with certain key factors, such as soil organic matter content, which can be estimated by measuring total C. Indeed, the rhizosphere soil of sorghum contained more total C than the bulk soil (Fig 2), and because soil organic matter contributes substantially to the retention of nutrient minerals via the provision of ionic functional groups as mineral adsorption sites, it is likely that the accumulation of organic matter in the rhizosphere was related to the enrichment of mineral nutrients. To examine this hypothesis, partial correlation coefficients were calculated for the parameters under the control of total C. Most of the significant correlations observed in Fig 4 disappeared in the analysis (Fig 5), which suggests that those correlations were indirectly derived from strong correlations between the parameters and total C. This is consistent with the idea that the accumulation of organic matter in the rhizosphere contributes to the enrichment of minerals. However, the origin of the organic matter accumulated in the rhizosphere remains unclear. If increases in total C were mainly due to the secretion of carbohydrates by plant roots or to the accumulation of cell wall debris, the accumulation of C-rich molecules should also increase the C/N ratio. However, the C/N ratios of the bulk and rhizosphere soils remained relatively similar, which suggests that the organic matter accumulated in the rhizosphere was the same substance as that found in the bulk soil. Thus, it is possible that the different levels (but not C/N ratios) of the organic matter in soil are owing to differences in microbe biomass.

**Fig 4.**
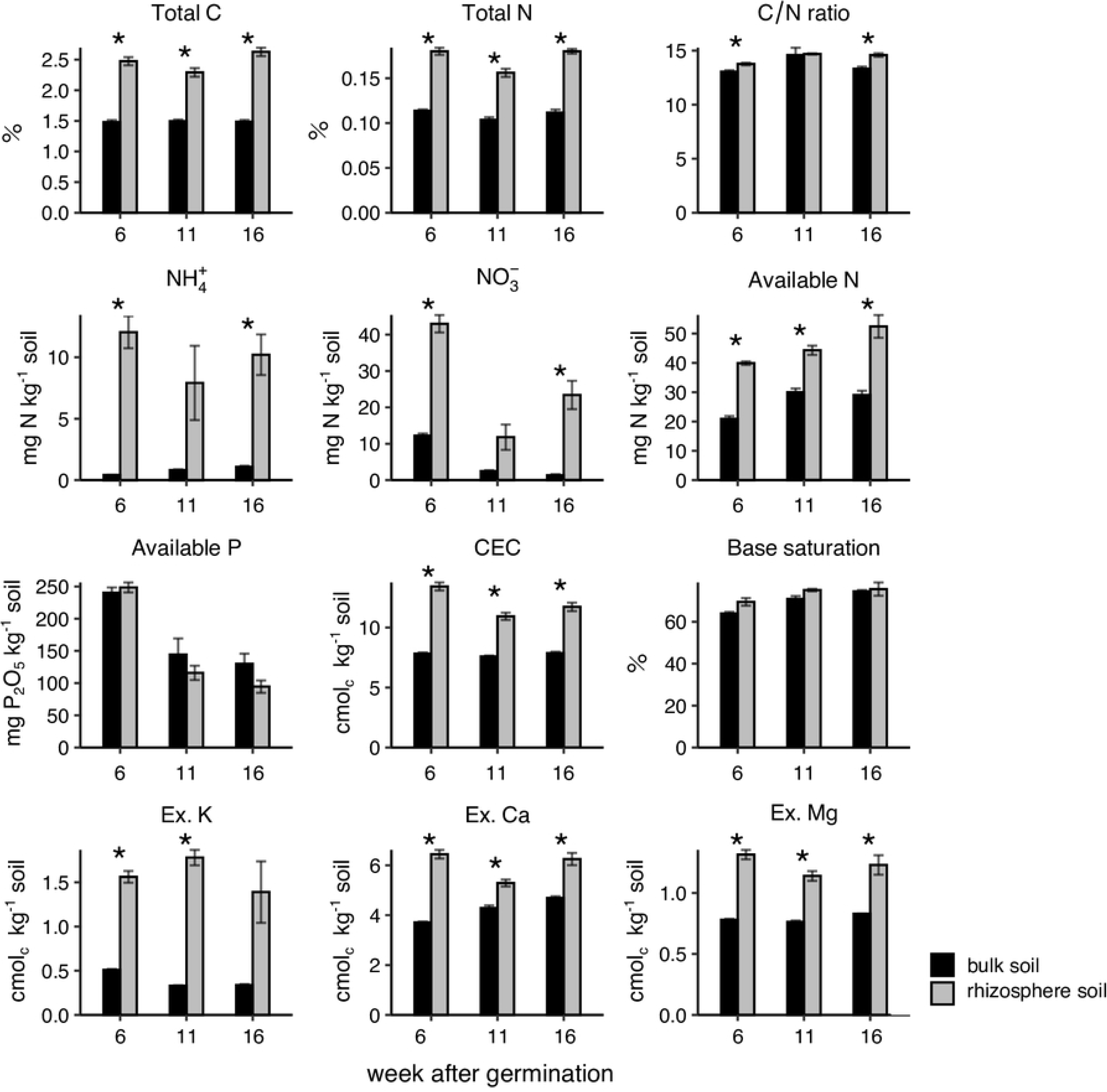
Correlations between the characteristics of container-grown sorghum soil. Values are Spearman correlation coefficients, and asterisks (* and **) indicate significant differences at *p* < 0.05 and 0.01, respectively. The cells that contain significant correlation values are colored according to the scale provided.

**Fig 5.**
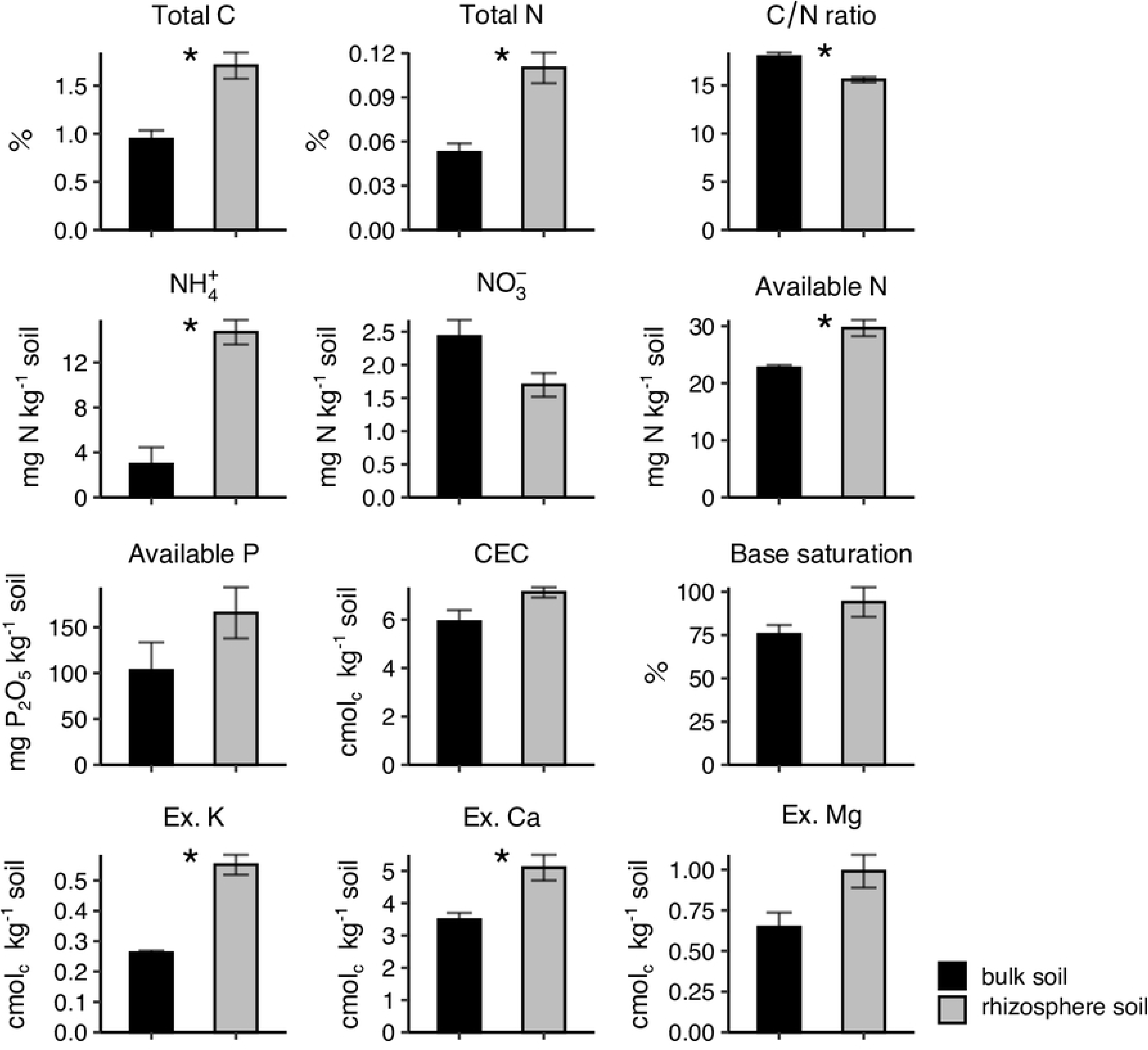
Partial correlation coefficients between the characteristics of container-grown sorghum soil. Values are Spearman correlation coefficients that were calculated after accounting for total soil C content. Asterisks (* and **) indicate significant differences at *p* < 0.05 and 0.01, respectively, and the cells that contain significant correlation values are colored according to the scale provided.

It is also possible that the enrichment of minerals could be related to the attraction of cations by negative charges on root surfaces. Plant cells are surrounded by cell walls that contain pectin, which is rich in carboxylic acid groups that impart fixed negative charges to cell surfaces. Thus, the effects of the negative charges on nutrient absorption by roots has been investigated using both cell walls and intact roots [25–27]. In addition, both the weathering of clay minerals and decomposition of soil organic matters can be enhanced in rhizosphere, through the action of plants and microorganisms. The accelerated weathering of clay minerals and mobilization of non-exchangeable K are thought to occur in the rhizospheres of Norway spruce and oak [28], and enhanced N mineralization has been reported to occur in rhizospheres of oat plants [29]. Such an increased rate of mineralization, together with the accumulation of source organic matter, might have contributed to the enrichment of NH_4_^+^ in the rhizosphere.

Among previous studies that have characterized the soil around roots, those that have analyzed root-adhering soils collected by mechanical isolation have reported that rhizosphere soils contain more minerals than bulk soils [5,6], whereas studies using rhizobox or similar methods report that the minerals are rather depleted in the soils around roots [8-10,12]. These discrepancies are probably due to differences in the distance between analyzed soils and root surfaces. Nutrient minerals are depleted around roots by plant uptake, but may actually be accumulated to higher levels at very close proximities to root surfaces. Possibly consistent with this hypothesis, a previous rhizobox study reported that exchangeable K in the soil associated with rape roots exhibited a decreasing gradient towards the root compartment but increased towards the root surface within 1 mm of the roots [11]. The authors suggested that the increase in K content could be due to the K content of root hairs or other plant tissues in the soil sample. However, such contamination does not account for the NH_4_^+^ enrichment observed in the present study (Figs 2 and 3) and reported by other previous studies [5,6] since any N accumulated in plant tissues should be either NO_3_^−^ or organic N compounds but not NH_4_^+^. Taken together, these results suggest that nutrient minerals accumulate around roots via multiple mechanisms, including the increased density of adsorption sites and enhanced generation from insoluble forms, but that such accumulation only occurs within very limited distances from the roots.

## Conclusion

In the present study, the nutrient mineral contents of the rhizosphere soils of herbaceous plants were analyzed using a set of small-scale protocols that were developed for this purpose. The nutrient mineral contents were generally greater in the rhizosphere than in bulk soil, especially in regards to NH_4_^+^ and exchangeable K. These results suggest that nutrient availability values that are estimated using bulk soil analysis are not representative of the nutrient availability of the rhizosphere, from which plants actually take up nutrients. More extensive surveys of rhizosphere mineral contents are necessary to address this issue. The small-scale protocols presented in the present study would be useful for performing such analyses.

## Acknowledgments

We are grateful to Dr. Ken Kawamoto (Saitama University) for the analysis of specific surface areas of soil particles, to EARTHNOTE Co., Ltd. for the gift of sorghum seeds, and to Drs. Naoki Moritsuka (Kyoto University) and Daisuke Shibata (Kazusa DNA Research Institute) for valuable discussion. This work was supported, in part, by CREST Grant (no. JPMJCR17O2) from Japan Science and Technology Agency (JST), and Science and Technology Research Partnership for Sustainable Development (SATREPS) Grant from Japan International Cooperation Agency (JICA) and JST.

